# Growth of Primary and Lateral Roots of Vicia faba L. in the Solution of Calcium Sulfate

**DOI:** 10.1101/007716

**Authors:** Michael Pyshnov

## Abstract

The presence of calcium sulfate in cultivating solution prevents bacterial contamination, browning/lignification and the death of the roots. There is no need for surface sterilization of seeds. No other ingredients, beside calcium sulfate, are needed for the healthy growth of the primary and lateral roots. There is no need to change the solution for several weeks when up to 60 lateral roots per seed can appear.

A modification of the two-stage growing technique where the seeds are first suspended in moist air over the cultivating solution to grow primary roots, and then, the primary roots are covered with the cultivating solution to grow lateral roots, was used.

The hypothesis is put forward that primary roots actually need water as a liquid to expel air, as the air is probably preventing the appearance of lateral roots.

## Introduction

The lateral roots (LR) of *V. faba* provide a desirable material for research on the large chromosomes of this species and other purposes. Without reviewing the current methods of seedlings cultivation, several examples can be given. Almost universally, the seeds undergo surface sterilization with potentially harmful chemicals, such as 15% Clorox, to avoid bacterial growth. Different substrates providing moisture to the roots are used. Vigorous aeration is sometimes recommended as a necessity. The number of ingredients in the cultivating solutions is ever increasing, the recommended concentrations widely differ.

The author has taken a view that, in the methods used, there is a substantial lack of appreciation of the different conditions that primary and LR roots need. Therefore, the procedure was modified to allow good separation of the first stage when primary roots need moist air, and the second stage when they are immersed in water. The use of calcium sulfate in the cultivating solution allowed to eliminate the potentially harmful surface sterilization. It was also found that other components of the cultivating solution can be disposed with.

## Materials and Methods

1. Initial experiments were carried out in plastic containers. Later, the glass containers were used in which the rim was filed down in four points to accommodate two stainless steel hangers holding the seeds, so that the lid could rest on the rim of the container (see Fig. 2).
2. Deionised and ozonised commercial water sold in plastic bottles was used.
3. Calcium sulfate dihydrate, 99% purity (Sigma-Aldrich Co, USA), was used to make 0.1% solution. It is referred to in the text as cultivating solution or simply solution.
4. Cultivating temperature was 19-20° C in the experiment on Fig. 1, but it was 20-25° C in the experiment on Fig. 2.
5. The relative humidity of air over the solution was measured as between 95% and 100%.
6. The experiments were carried out in dark box.
7. The procedure with the seeds was as follows:

1. Small seeds and seeds with damaged coat were rejected.
2. The hole was drilled in seeds for the hangers.
3. Washing the seeds in the solution for 1/2 hour several times.
4. Soaking seeds for 24 hours in the solution.
5. Washing.
6. Peeling the coat.
7. Washing.
8. Hanging seeds on wires and placing in the containers.
9. When primary roots attained sufficient length, the cultivating solution was added to cover the roots.
10. The shoots were cut off when they reached 2-3 cm.
8. Photographs were taken twice a day with digital camera. The obtained jpeg images at 180 dpi were converted to pdf format, cropped and pasted to Word and finally - all converted to pdf. No “photoshopping” was used.

## Results

The author has tested various compounds, substrates and techniques for over a year. This resulted in the finding that calcium sulfate stops bacterial contamination and browning of the roots practically completely. It became the only ingredient used further in the cultivating solution. (Calcium sulfate, gypsum, has been used to prevent root rot of various bacterial origins for at least the last forty years.)

**Fig. 1.**
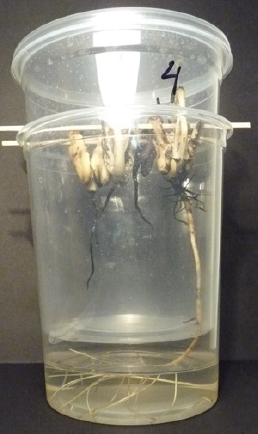
Four seeds suspended on wooden sticks, were cultivated in plastic containers. They were sprayed with the solution two or three times a day. Such spraying leads to severe browning, and, if lateral roots appear, they soon also die. But in a rare event here, one primary root survived and continued to grow, not producing lateral roots. It went through a small hole in the bottom of the upper container to the solution below, reaching 11 cm in length. At this point, it started producing lateral roots in the part immersed in the solution. This photo was taken 20 days after soaking the seeds.

The observations have shown that moist filter paper is providing irregular contacts of roots with water, with air, and with the paper, and this, apparently, was causing highly irregular root behaviour. Therefore, the water providing substrate was disposed with; the roots were suspended at both stages, first in moist air and then in the solution.

The early experiment below on Fig. 1. showed that primary root can grow in moist air, but the lateral roots appear only when it is immersed in water.

The primary root in Fig. 1. was only slightly affected by browning, but in the 7 days that it was growing in moist air, or at any time during the 20 days, it did not produce LR in air. The multiple LR, however, appeared after 2 days under the water. This cannot be explained solely by the effect of browning or the absence of such.

Primary roots growing in air were capable of producing LR when they attained the length of some 3 cm. Some observations showed that shorter primary roots suspended close to the water below, would make 90° turn, as if avoiding contact with water, and they continued to grow horizontally in air for some time. Also, the dry seeds, washed and peeled in the cultivating solution, and left in the solution for many days, swell and increase in size several times, but the primary root never grows, let alone produces LR: it needs air. Again, it grows perfectly well in water after producing LR.

The Fig. 2. on the next page shows the consecutive stages of roots growth after the seeds were washed, soaked for 24 hours in the solution and peeled prior to the incubation in moist air over the solution (see Methods).

**Fig. 2.**
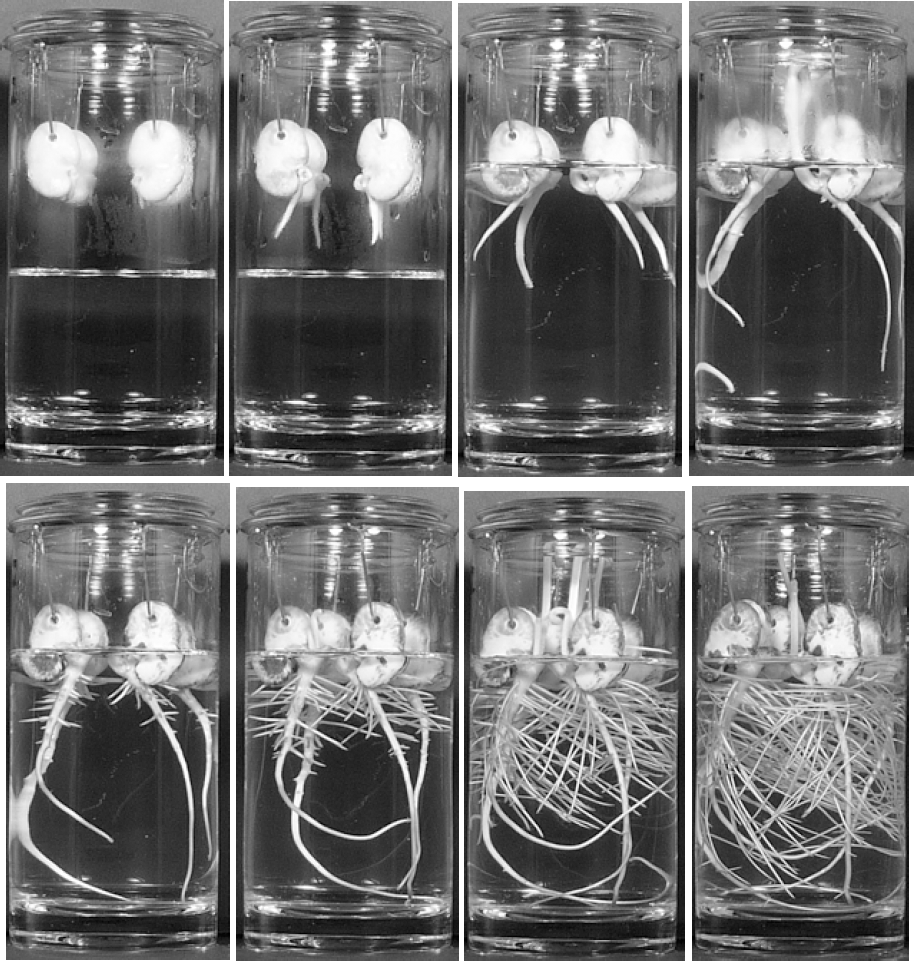
The time, days after soaking, from left to right, was as follows: Top: 2, 3, 4, 6 days. Bottom: 7, 8, 10, 15 days. The solution was added immediately after 3-days photo was taken. Barely visible pimples of emerging LR appeared after 5 1/2 days. There are considerable spots of browning on the cotyls, but all roots are completely free from it. On some photos the dew on the glass obscures the image. Diameter of the glass container is 9 cm.

## Discussion

Immersion of seedlings in water to grow LR has been used for a very long time [1]. However, they were usually first germinated on moist paper or cotton and often even never transferred to water. The current papers and reviews dealing with the subject, often do not mention the central fact - that the primary root which grows in moist air, must be immersed in water to grow LR. The results of the present work are in agreement with the part of the recently published results that deals with the “patterning” of LR by water in *Arabidopsis* [2]. However, it seems that the better interpretation can be that, simply, lateral roots appear in water, but not in the air.

### The hypothesis

The need for the primary root to be covered with water to produce LR, even when the humidity is high, is a most puzzling fact. It is very doubtful that water as a chemical component is unavailable to the root in such air. So, the author came to the hypothesis that primary root actually needs water as a liquid to expel air, as the air is probably preventing the appearance of LR.

### Calcium sulfate

The use of calcium sulfate helped greatly to see what is actually needed by the roots. It is not known if calcium sulfate had nutritional value here, however, changing the solution at later stages had no effect on roots growth, which also suggests that no great amounts of poisonous material were accumulating. Apparently, in these experiments, root nutrition is provided by the usual source - the cotyledons. It is also possible that residual bacteria are participating. And, it cannot be excluded that there occurs a crystallisation of calcium sulfate on the primary roots when they grow in moist air, which, generally speaking, can influence the process leading to the appearance of LR.

## References

1. Trosko, J. E. and Wolff, S. (1965) Strandedness of Vicia faba chromosomes as revealed by enzyme digestion studies. J. Cell Biol. 26, 125–135.

2. Yun Bao, et al. (2014) Plant roots use a patterning mechanism to position lateral root branches toward available water. Proc. Nat. Acad. Sc., 111(25): 9319–9324.

